# Hand Grip Strength, Cognitive Function and the Role of Cognitive Reserve: Results from a Sample of Community Dwelling Elderly in China

**DOI:** 10.1101/2019.12.18.881037

**Authors:** Rong Wei, Kai-yong Liu, Fang-biao Tao, Pei-ru Xu, Bei-jing Cheng, Liang Sun, Qu-nan Wang, Qiang-wei Feng, Xiu-de Li, Lin-sheng Yang

**Affiliations:** School of Public Health, Department of Nutrition and Food Hygiene, Anhui Medical University, Anhui Hefei, 230032, China; School of Public Health, Department of Epidemiology and Health Statistics, Anhui Medical University, Anhui Hefei, 230032, China; School of Health Services Management, Anhui Medical University, Anhui Hefei, 230032, China; Anhui Provincial Key Laboratory of Population Health and Aristogenics, Anhui Hefei, 230032, China; School of Public Health, Department of toxicology, Anhui Medical University, Anhui Hefei, 230032, China; Fuyang Center for Disease Control and Prevention, Anhui Fuyang, 236000, China; Lu’an Center for Disease Control and Prevention, Anhui Lu’an, 237008, China

**Author notes:** The first two authors contributed equally to this paper. **Corresponding author:** Fang-biao TAO, School of Health Services Management, Anhui Medical University, Anhui Provincial Key Laboratory of Population Health and Aristogenics, Meishan Road 81 Hefei, Anhui, 230032, China; Fax: none. Lin-sheng YANG, Department of Epidemiology and Health Statistics, Anhui Medical University, Meishan Road 81 Hefei, Anhui, 230032, China; Fax: none.

**Keywords:** cognitive function, cognitive reserve, hand grip strength, older adults

## Abstract

**Objectives:** To examine the association between hand grip strength (HGS) and cognitive function, and the potentially moderating effects of cognitive reserve on this relationship using a sample of community dwelling elderly in China.

**Methods:** The subjects included 1291 community-dwelling elderly aged 60 or over and without dementia who participated in the baseline survey of an elderly cohort in Anhui province, China. Cognitive function was assessed using Mini-Mental State Examination (MMSE) and HGS was measured using an electronic grip strength dynamometer. The education (EDU) in early life, cognitive level of the job (CLJ) in middle age, cognitive leisure activities (CLA) in late life, and other covariates were collected through a face-to-face interview and physical examination.

**Results:** The differences in MMSE scores across tertiles of HGS were significant (MMSE scores across tertiles of HGS: 20.26±7.02 vs 22.83±5.99 vs 24.76±6.36, *F*=62.05, *P*<0.001). After adjustment for covariates, the lower tertiles of HGS was related to lower MMSE scores when compared to the upper tertiles of HGS (*β*=*β* [*95%CI*]: −2.02[−2.87~−1.17], *P*<0.001). However, no significant association existed between the intermediate tertiles of HGS and lower MMSE scores (*β*=*β* [*95%CI*]: −0.28[−1.05~0.50], *P*=0.483). Moderation analyses revealed that the correlation between the lower tertiles of HGS and decreased MMSE scores was less pronounced in middle EDU (*β*=*β* [*95%CI*]: −1.62[−3.22~−0.02], *P*=0.047), and in middle CLJ (*β*=*β* [*95%CI*]: −2.17[−3.31~−1.24], *P*<0.0001) than in low EDU (*β*=*β* [*95%CI*]: −2.46[−3.80~−1.12], *P*<0.0001), and in low CLJ (*β*=*β* [*95%CI*]: −3.72[−6.92~−0.53], *P*=0.023). Furthermore, this relationship was not significant among the elderly with high EDU or the high CLJ.

**Conclusions:** The lower HGS is associated with poor cognitive function in older age, and cognitive reserve may attenuate or eliminate the relationship of lower HGS with cognitive function.

## 1. Introduction

Dementia is one of the greatest contemporary health and social care challenges. Globally, about 47 million people were living with dementia in 2015, and the number is projected to almost double every 20 years [1,2]. Due to the rapidly aging population in China, the challenge posed by dementia or cognitive impairment is much greater than that reported from developed countries [1]. No effective treatments for dementia have been so far obtained, thus strategies to identify modifiable risk factors for preventing or delaying the onset of dementia are urgently warranted.

Among modified factors for dementia or cognitive impairment in old age, hand grip strength (HGS) is extensively investigated. Numerous studies [3–5] have shown that reduced HGS over time may be served as a predictor of cognitive loss with advancing age. For example, Taekema and colleagues [4] have discovered that lower HGS at baseline could predict an accelerated decline in MMSE scores during the 4-year follow-up period. Buchman and colleagues [5] have reported that in a sample of 877 older adults followed for 5.7 years, an annual decrease of 1 1b in HGS was associated with a 9% increase in the risk for Alzheimer’s disease. Several hypotheses have been proposed to explain the relationship between lower HGS and cognitive impairment. Stronger HGS may reflect the integrity of neuromuscular system and higher resistance to oxidative stress and inflammation, which may contribute to preservation of cognitive function [6,7]. Nonetheless, a few inconsistent results have also been obtained in other cohort studies. For example, Sternäng et al [8] found the association between reduced HGS and decline in cognitive function during follow-up period varied by age. Reduced HGS predicted decline in verbal ability, spatial ability, processing speed, and memory only in individuals over the age of 65 rather than in those under age 65. Atkinson et al [9] have reported that decline in HGS did not predict decline in cognitive function over time in the Women’s Health Initiative Memory Study. Similar results was also reported by Gray [10] that in a large American sample with being predominantly white and well educated, lower HGS was not associated with an increased risk of developing dementia. These inconsistent results imply that some other factors may modify the relationship between HGS and cognitive function.

Cognitive reserve is considered as a buffer for the relationship of pathology with cognitive decline [11–13]. The concept of cognitive reserve provides an explanation for individual differences in processing cognitive tasks which allow some to cope better than others with brain pathology [11–13]. Cognitive reserve may be based on more efficient utilization of brain networks or of enhanced ability to recruit alternate brain networks as needed [11–13]. These mechanisms may contribute to the adaptation of brain activity and improvement of cognitive performance in individuals when task difficulty is increased. Epidemiological studies have found that stimulating activities at different stages of life (i.e. educational and occupational attainment, and cognitive leisure activity) could contribute to increasing cognitive reserve and attenuate age-related cognitive impairment [14–22]. For example, Galioto RM et, al. have found a significant interaction between body mass index (BMI) and cognitive reserve on attention/executive function and memory, suggesting that cognitive reserve may attenuate obesity-related cognitive impairment [17]. Hence, it seems an appealing question whether or not cognitive reserve would attenuate the association of HGS with cognitive function. A recent study found that the relation of cognitive reserve to MMSE scores varied as a function of grip strength in a sample of community-dwelling older adults in Brazil [18]. To our knowledge, no study has focused on the role of cognitive reserve on the association of HGS with cognitive function, which is of great importance given that cognitive reserve may be a early target in preventing the onset of dementia.

The current study aims to expand previous studies by testing the association between HGS and cognitive function with a strict control of confounding factors, and to examine the moderating effect of cognitive reserve on this relation using a sample of Chinese older adults. We hypothesized that lower HGS would be associated with lower MMSE scores, and this association would be less pronounced in higher level of cognitive reserve.

## 2. Methods

### 2.1 Study design and participants

Data of this study came from the baseline survey of an elderly cohort: *Elderly Health and Modified Factors*, which was launched in lu ‘an city and Fuyang city, Anhui province, China, from June to September in 2016. This cohort was built by Department of Public Health of Anhui Medical University and local Center for Disease Control and Prevention. Firstly, two counties were randomly selected from the lu ‘an city and one county was randomly selected from the Fuyang city. Secondly, one community in each county was randomly selected. Finally, 500 community-dwelling elderly aged 60 or above in each community were invited to attend this survey.

Inclusion criteria for participants were as follows: no symptoms of dementia, having been living in one of the three geographic areas of Anhui province for at least half a year, and being conscious and acting freely. Exclusion criteria for participants were as follows: having symptoms of Parkinson’s disease, having been suffering from stroke, unwillingness to cooperate with investigators. Individuals were volunteers recruited via advertisements by Center for Disease Control and Prevention and community hospitals. Community doctors made telephone appointments with the elderly one day in advance considering the response rate. Eligible individuals were asked to complete an extensive face-to-face interview and physical examination conducted in the local community hospitals which lasted for 2~3 hours after signing a written informed consent form. All protocols of this study were reviewed and approved by the Ethics Committee of Anhui Medical University.

### 2.2 Cognitive function

Cognitive function was examined using the Mini-Mental State Examination (MMSE) [23] which includes multiple cognitive abilities (orientation, memory, counting backwards, and language). MMSE has been widespread used to assess cognitive status of the elderly and its scores range from 0 to 30, with lower scores indicating poorer cognitive function. Community-dwelling elderly were asked to answer the questions and respond to the investigator’s requests immediately. In this study, the Cronbach’s alpha reliability coefficient was 0.79.

### 2.3 Hand grip strength (HGS)

We measured HGS to the nearest kilogram using a hand dynamometer (JH-1881, China). The participants stood in an upright position with the measured arm unsupported. The dynamometer was parallel to the body, which slightly away from the body. Participants had to squeeze firmly and gradually and build quickly up to maximum force. After the procedure was explained to each individual, three trials of HGS were performed for each hand, and the average of three trials was calculated and recorded. Three tests were performed on the same day with one minute gap between every two tests. The dominant grip was used for scoring purpose which was the maximum of two mean values.

HGS was divided into tertiles by gender considering the great difference between males and females in HGS. For males, lower, intermediate, and upper tertiles were 3.5-26.2 kg, 26.2-33.4 kg, 33.4-58.4 kg, respectively. For females, lower, intermediate, and upper tertiles were 2.0-17.0 kg, 17.0-22.0 kg, 22.0-42.9 kg, respectively.

### 2.4 Markers of Cognitive Reserve

#### 2.4.1 Education (EDU)

Participants were asked to indicate their highest educational level, with a five-level option as follows: (1) illiteracy level, (2) primary school level, (3) junior high school level, (4) technical secondary school or apprenticeship graduation or senior high school level, (5) superior vocational college degree or university degree. Considering the limited sample size, we divided educational level into three EDU levels: low (option 1), middle (option 2), and high (option 3 through 5) [14, 17, 18, 20].

#### 2.4.2 Cognitive Level of the job (CLJ)

Participants were asked to identify the principal profession they had practiced during their working life, which we thereafter classified into 3 CLJ groups by the degree of intellectual involvement at work: (1) low (e.g. factory work, plumbing, carpentry, farming, etc.), (2) middle (e.g. production personnel, the driver, etc), (3) high (e.g. craftsman, clerk, trader, teacher, clerical work, medical practice, etc) [16–18, 20].

#### 2.4.3 Cognitive leisure activity (CLA)

Participants were asked to report the average time per day of cognitively stimulating leisure activities in the past two months such as playing cards, scrabble, using computer, or reading, etc. We divided CLA into three groups according to the length of time: low (<1 hour per day), middle (1~3 hours per day), high (*>*3 hours per day) [15, 17–19, 21, 22].

### 2.5 Covariates

Sociodemographic factors consisted of sex, age (5-year bands), marital status (divorced or unmarried, married, and widowed), and residence (urban and rural).

Health behavioral factors were assessed by questionnaire and included smoking status (yes and no), alcohol consumption (yes and no), and physical exercise per day (yes and no).

Traditional cardiovascular (CV) risk factors included body mass index (BMI) (categorised as <18.5, 18.5-24.0, 24.0~28.0, and ⩾ 28.0 kg/m^2^)[24], total level of cholesterol (divided into tertiles: <5.2, 5.2~6.0, and >6.0 mg/dl), triglycerides (divided into tertiles: <1.2, 1.2~1.7, and >1.7 mg/dl), hypertension (determined by systolic blood pressure ⩾ 140 mm Hg, diastolic blood pressure ⩾ 90 mm Hg, reported doctor diagnosed hypertension, use of antihypertensive drugs, or hospital records), diabetes (determined by fasting blood glucose ⩾ 7.0 mmol/L, reported doctor diagnosed diabetes, use of antidiabetic drugs, or hospital records), coronary heart disease identified using hospital medical record or self reported use of drugs for coronary heart disease, and depressive symptoms assessed by 30-item Geriatric Depression Scale[25].

### 2.6 Statistical Analyses

Firstly, contingency table was used to describe the distribution of teritles of HGS across sociodemographic factors, health behavioral factors, CV risk factors, EDU, CLJ, and CLA. Secondly, a t-test or ANOVA was used to assess the differences of MMSE scores between different variables such as teritles of HGS, demographic variables, health behavioral factors, CV risk factors, EDU, CLJ, and CLA. Thirdly, multiple linear regression model was used to inspect the relationship between cognitive function and teritles of HGS after controlling for sociodemographic variables, health behavioral factors, CV risk factors, EDU, CLJ, and CLA. Then three interaction terms (HGS and EDU, HGS and CLJ, HGS and CLA) were included into the multivariate model, respectively. If a statistical significance existed for any one of the interaction terms, the relation of HGS to MMSE scores stratified by EDU, CLJ, or CLA would be shown, respectively.

All analyses were performed using SPSS software version 23.0. Two-sided *P* values were used with an α level of 0.05 for statistical significance.

## 3. Results

### 3.1 Descriptive Statistics

A total of 1,500 persons accepted to take part in this survey. After following further exclusions of 80 participants who had missing cognitive status information or did not provided complete scale data and 150 participants who had missing HGS information or did not involved in the measurement of HGS, the core analytic sample size was 1291. No significant differences were observed in age (mean age, 70.9 and 70.7 years) and ratio of males to females (55% vs 53%) between participants included and those excluded (all p-values > 0.05).

Of 1291 persons, mean age was 71 years (SD = 6.4, range: 60-94). Males accounted for 44.6% (n=576). Mean MMSE score was 22.56 (SD = 6.36; range 10-30). About 65.1% participants lived in rural residence. The proportion of participants being divorced or never-married, widowed, and married was 5.0%, 20.2%, and 73.8%, respectively.

Table 1 showed that the differences were statistically significant in distribution of HGS tertiles across age, marital status, smoking, alcohol consumption, triglycerides, coronary heart disease, depressive symptoms, EDU, CLJ, and CLA. Table 2 showed that the differences of MMSE scores were statistically significant across HGS, age, sex, residence, marital status, smoking, alcohol consumption, physical exercise, BMI, triglycerides, depressive symptoms, EDU, CLJ, CLA.

**Table 1.** Distribution of HGS teritles across different variables

| HGS teritles |
| --- |

| Variables | N | lower<br>N=435 | intermediate<br>N=431 | upper<br>N=425 | $F/\chi^2$ | $P$<br>value |
| --- | --- | --- | --- | --- | --- | --- |
| Sociodemographic Factors |  |  |  |  |  |  |
| Age* (years old) |  |  |  |  |  |  |
| 60-65 | 346 | 79(22.8) | 120(34.7) | 147(42.5) | 85.14 | <0.001 |
| 66-70 | 361 | 99(27.4) | 126(34.9) | 136(37.7) |  |  |
| 71-75 | 278 | 97(34.9) | 99(35.6) | 82(29.5) |  |  |
| 76-94 | 300 | 160(53.3) | 83(27.7) | 57(19.0) |  |  |
| Sex |  |  |  |  |  |  |
| males | 576 | 194(33.7) | 192(33.3) | 190(33.0) | 0.034 | 0.983 |
| females | 715 | 241(33.7) | 239(33.3) | 235(33.0) |  |  |
| Residence |  |  |  |  |  |  |
| rural | 840 | 280(33.3) | 276(32.9) | 284(33.8) | 0.87 | 0.647 |
| urban | 451 | 155(34.4) | 155(34.4) | 141(31.3) |  |  |
| Marital status* |  |  |  |  |  |  |
| Divorced/unmarried | 65 | 28(43.1) | 23(35.4) | 14(21.5) | 88.88 | <0.001 |
| widowed | 261 | 142(54.4) | 82(31.4) | 37(14.2) |  |  |
| married | 953 | 258(27.1) | 324(34.0) | 371(38.9) |  |  |
| Health Behavioral Factors |  |  |  |  |  |  |
| Smoking |  |  |  |  |  |  |
| no | 892 | 294(33.0) | 322(36.1) | 276(30.9) | 9.39 | 0.009 |
| yes | 399 | 141(35.3) | 109(27.3) | 149(37.3) |  |  |
| Alcohol consumption |  |  |  |  |  |  |
| no | 832 | 324(38.9) | 317(38.1) | 191(23.0) | 99.08 | <0.001 |
| yes | 459 | 111(24.2) | 114(24.8) | 234(51.0) |  |  |
| Physical exercise |  |  |  |  |  |  |
| no | 929 | 339(36.5) | 317(34.1) | 310(33.4) | 3.66 | 0.160 |
| yes | 362 | 96(26.5) | 114(31.5) | 115(31.8) |  |  |
| CV risk factors |  |  |  |  |  |  |
| BMI* (kg/m<sup>2</sup>) |  |  |  |  |  |  |
| <18.5 | 61 | 26(42.6) | 20(32.8) | 15(24.6) | 6.21 | 0.400 |
| 18.5~24.0 | 554 | 186(33.6) | 190(34.3) | 178(32.1) |  |  |
| 24.0~28.0 | 458 | 140(30.6) | 155(33.8) | 163(35.6) |  |  |
| >28.0 | 190 | 70(36.8) | 59(31.1) | 61(32.1) |  |  |
| Total cholesterol (mg/dl) |  |  |  |  |  |  |
| <5.2 | 485 | 158(32.6) | 145(29.9) | 182(37.5) | 8.50 | 0.075 |
| 5.2~6.0 | 354 | 118(33.3) | 129(36.4) | 107(30.2) |  |  |
| >6.0 | 452 | 159(35.2) | 157(34.7) | 136(30.1) |  |  |
| Triglycerides (mg/dl) |  |  |  |  |  |  |
| <1.2 | 484 | 154(31.8) | 154(31.8) | 176(36.4) | 9.51 | 0.050 |
| 1.2~1.7 | 349 | 119(34.1) | 108(30.9) | 122(35.0) |  |  |
| >1.7 | 458 | 162(35.4) | 169(36.9) | 127(27.7) |  |  |
| **Hypertension** |  |  |  |  |  |  |
| no | 393 | 134(34.1) | 133(33.8) | 126(32.1) | 0.19 | 0.910 |
| yes | 898 | 301(33.5) | 298(33.2) | 299(33.3) |  |  |
| **Diabetes\*** |  |  |  |  |  |  |
| no | 1015 | 344(33.9) | 331(32.6) | 340(33.5) | 1.44 | 0.487 |
| yes | 275 | 90(32.7) | 100(36.4) | 85(30.9) |  |  |
| **Coronary heart disease\*** |  |  |  |  |  |  |
| no | 1103 | 349(31.6) | 376(34.1) | 378(34.3) | 11.00 | 0.004 |
| yes | 162 | 72(44.4) | 49(30.2) | 41(25.3) |  |  |
| **Depressive symptoms\*** |  |  |  |  |  |  |
| no | 858 | 251(29.3) | 290(33.8) | 317(36.9) | 29.05 | <0.001 |
| yes | 407 | 176(43.2) | 131(32.2) | 100(24.6) |  |  |
| **Markers of Cognitive Reserve** |  |  |  |  |  |  |
| **EDU** |  |  |  |  |  |  |
| low | 641 | 273(42.6) | 225(35.1) | 143(22.3) | 75.64 | <0.001 |
| middle | 302 | 77(25.5) | 100(33.1) | 125(41.4) |  |  |
| high | 348 | 85(24.4) | 106(30.5) | 157(45.1) |  |  |
| **CLJ** |  |  |  |  |  |  |
| low | 876 | 317(36.2) | 291(33.2) | 268(30.6) | 9.56 | 0.049 |
| middle | 96 | 28(29.2) | 32(33.3) | 36(37.5) |  |  |
| high | 319 | 90(28.2) | 108(33.9) | 121(37.9) |  |  |
| **CLA (hours per day)** |  |  |  |  |  |  |
| low | 338 | 127(37.6) | 113(33.4) | 98(29.0) | 6.90 | 0.141 |
| middle | 710 | 219(30.8) | 242(34.1) | 249(35.1) |  |  |
| high | 243 | 89(36.6) | 76(31.3) | 78(32.1) |  |  |

**Abbreviations:** \*, data missing; MMSE, Mini-mental state examination; HGS, hand grip strength (kg); BMI, body mass index (kg/m<sup>2</sup>); CV risk factors, cardiovascular risk factors; EDU, education; CLJ, cognitive level of the job; CLA, cognitive leisure activity.

**Table 2.** Comparison of MMSE across different variables

| Variables | N | MMSE( $\bar{x} \pm s$) | F/t | P |
| --- | --- | --- | --- | --- |
| **HGS\* (kg)** |  |  |  |  |
| lower | 435 | 20.26±7.02 | 62.05 | <0.001 |
| intermediate | 431 | 22.83±5.99 |  |  |
| upper | 425 | 24.76±6.36 |  |  |

| **Sociodemographic Factors** |  |  |  |  |
| --- | --- | --- | --- | --- |
| **Age\*** (years old) |  |  |  |  |
| 60-65 | 346 | 23.22±5.49 | 14.45 | <0.001 |
| 66-70 | 361 | 23.38±5.60 |  |  |
| 71-75 | 278 | 23.00±6.37 |  |  |
| 76-94 | 300 | 20.52±7.59 |  |  |
| **Sex** |  |  |  |  |
| males | 576 | 23.93±6.03 | 49.48 | <0.001 |
| females | 715 | 21.47±6.42 |  |  |
| **Residence** |  |  |  |  |
| rural | 840 | 20.82±6.35 | 210.40 | <0.001 |
| city | 451 | 25.82±4.97 |  |  |
| **Marital status\*** |  |  |  |  |
| divorced or never-married | 65 | 18.09±7.20 | 26.24 | <0.001 |
| widowed | 261 | 21.48±7.18 |  |  |
| married | 953 | 23.22±5.84 |  |  |
| **Health Behavioral Factors** |  |  |  |  |
| **Smoking** |  |  |  |  |
| no | 892 | 23.00±6.46 | 12.15 | 0.001 |
| yes | 399 | 21.69±6.43 |  |  |
| **Alcohol consumption** |  |  |  |  |
| no | 833 | 21.75±6.67 | 44.20 | <0.001 |
| yes | 458 | 24.14±5.83 |  |  |
| **Physical exercise** |  |  |  |  |
| no | 929 | 21.62±6.74 | 86.00 | <0.001 |
| yes | 362 | 25.12±4.95 |  |  |
| **CV risk factors** |  |  |  |  |
| **BMI\* (kg/m<sup>2</sup>)** |  |  |  |  |
| <18.5 | 61 | 19.59±7.14 | 8.28 | <0.001 |
| 18.5~24.0 | 554 | 22.11±6.56 |  |  |
| 24.0~28.0 | 458 | 23.15±6.09 |  |  |
| >28.0 | 190 | 23.55±5.93 |  |  |
| **Total cholesterol (mg/dl)** |  |  |  |  |
| <5.200 | 485 | 22.92±6.26 | 1.19 | 0.304 |
| 5.201~6.000 | 354 | 22.39±6.78 |  |  |
| >6.000 | 452 | 22.32±6.12 |  |  |
| **Triglycerides (mg/dl)** |  |  |  |  |
| <1.200 | 484 | 21.38±6.55 | 16.43 | <0.001 |
| 1.201~1.7161 | 349 | 22.69±6.26 |  |  |
| >1.7162 | 458 | 23.72±6.02 |  |  |
| **Hypertension** |  |  |  |  |
| no | 393 | 22.34±6.19 | 0.73 | 0.393 |
| yes | 898 | 22.66±6.43 |  |  |
| **Diabetes\*** |  |  |  |  |
| no | 1015 | 22.47±6.33 | 1.63 | 0.202 |
| yes | 275 | 23.01±6.33 |  |  |
| **Coronary heart disease\*** |  |  |  |  |
| no | 1103 | 22.47±6.39 | 3.65 | 0.056 |
| yes | 162 | 23.49±5.89 |  |  |
| **Depressive symptoms\*** |  |  |  |  |
| no | 858 | 24.10±5.54 | 152.27 | <0.001 |
| yes | 407 | 19.79±6.30 |  |  |
| **Markers of Cognitive Reserve** |  |  |  |  |
| **EDU** |  |  |  |  |
| low | 641 | 19.48±6.34 | 212.48 | <0.001 |
| middle | 302 | 24.40±5.07 |  |  |
| high | 348 | 26.66±4.08 |  |  |
| **CLJ** |  |  |  |  |
| low | 876 | 21.01±6.43 | 98.99 | <0.001 |
| middle | 96 | 24.09±5.97 |  |  |
| high | 319 | 26.37±4.24 |  |  |
| **CLA (hours per day)** |  |  |  |  |
| low | 338 | 22.37±6.45 | 11.30 | <0.001 |
| middle | 710 | 22.07±6.04 |  |  |
| high | 243 | 24.28±6.87 |  |  |

**Abbreviations:** \*, data missing; MMSE, Mini-mental state examination; HGS, hand grip strength (kg); BMI, body mass index (kg/m<sup>2</sup>); CV risk factors, cardiovascular risk factors; EDU, education; CLJ, cognitive level of the job; CLA, cognitive leisure activity.

### 3.2 Association of HGS and cognitive reserve with cognitive function

Compared to the upper tertiles of HGS, the lower (*β*=*β* [95%CI]: −0.70[−4.49~-2.91], *P*<0.001) and intermediate tertiles of HGS (*β*=*β* [95%CI]: −1.02[−1.82~−0.23], *P*=0.012) were negatively associated with MMSE scores in single-factor model. When adjustment for covariates including sociodemographic factors, health behavioral factors, CV risk factors, EDU, CLJ, and CLA, a statistically significant association between the lower tertiles of HGS and decreased MMSE scores still existed (*β*=*β* [95%CI]: −2.02[−2.87~−1.17], *P<*0.001). By contrast, the statistical significance disappeared in the association between the intermediate tertiles and decreased MMSE scores (*β*=*β* [95%CI]: −0.28[−1.05~0.50], *P=*0.483) (see table 3).

**Table 3.** The association between HGS and cognitive function in community dwelling elderly in China

| | $\beta$ | 95%CI | t value | P |
| --- | --- | --- | --- | --- |
| **Single-Factor Model** |  |  |  |  |
| **HGS Tertiles (kg)** |  |  |  |  |
| lower | -0.70 | -4.49~-2.91 | -9.12 | <0.001 |
| intermediate | -1.02 | -1.82~-0.23 | -2.53 | 0.012 |
| upper | ref. |  |  |  |
| **Multivariate model\*** |  |  |  |  |
| **HGS Tertiles<sup>a</sup> (kg)** |  |  |  |  |
| lower | -2.02 | -2.87~-1.17 | -4.67 | <0.001 |
| intermediate | -0.28 | -1.05~0.50 | -0.70 | 0.483 |
| upper | ref. |  |  |  |
| **EDU** |  |  |  |  |
| low | -4.57 | -5.38~-3.64 | -10.17 | <0.001 |
| middle | -1.11 | -1.97~-0.25 | -2.53 | 0.012 |
| high | ref. |  |  |  |
| **CLJ** |  |  |  |  |
| low | -1.01 | -1.96~-0.07 | -2.10 | 0.036 |
| middle | -0.83 | -2.06~0.41 | -1.32 | 0.189 |
| high | ref. |  |  |  |
| **CLA** |  |  |  |  |
| low | -0.32 | -1.19~0.55 | -0.72 | 0.470 |
| middle | -0.66 | -1.46~0.14 | -1.62 | 0.105 |

| high | ref. |
| --- | --- |

**Abbreviations:** *CI*, confidence interval; HGS, hand grip strength (kg); EDU, education; CLJ, cognitive level of the job; CLA, cognitive leisure activity; ref., reference.
\* Adjustment for age, sex, residence, marital status, smoking, alcohol consumption, physical exercise, BMI, total cholesterol, triglycerides, hypertension, diabetes, coronary heart disease, depressive symptoms, EDU, CLJ, and CLA.

Multivariate model also showed that lower MMSE scores were found in elderly with low level of EDU (*β*=*β* [95%CI]: −4.57[−5.38~−3.64], *P*<0.001), the middle level of EDU (*β*=*β* [95%CI]: −1.11[−1.97~-0.25], *P*=0.012), low level of CLJ (*β*=*β* [95%CI]: −1.01[−1.96~-0.07], *P*=0.036), compared to the high level of EDU or CLJ. However, no statistical significance was found in difference of MMSE scores between elderly with middle level of CLJ and those with high level of CLJ (*β*=*β* [95%CI]: −0.83[−2.06~0.41], *P=*0.189). No statistical significance was also found in difference of MMSE scores across levels of CLA (see table 3).

### 3.3 Moderation analyses

Two interaction terms were statistically significant (HGS×EDU: *β*=−0.729, *p*<0.001; HGS×CLJ: *β*=−0.609, *p*=0.004). Stratification analyses were then conducted, shown in table 4. The coefficients of the correlation between lower tertiles of HGS and poor cognitive function in individuals with low, middle, and high level of EDU were −2.46(95%*CI*:-3.80~-1.12), −1.62(95%*CI*:−3.22~−0.02), and −0.96(95%*CI*:−2.39~0.47), respectively.

**Table 4.** The association between HGS and cognitive function: stratification by EDU and CLJ

| | $\beta$ | 95%CI | $t$ | $P$ |
| --- | --- | --- | --- | --- |
| **Stratification by EDU\*** |  |  |  |  |
| **low** |  |  |  |  |
| HGS Tertiles (kg) |  |  |  |  |
| lower | -2.46 | -3.80~-1.12 | -3.61 | <0.001 |
| intermediate | -0.22 | -1.51~1.06 | -0.34 | 0.733 |
| upper | ref. |  |  |  |
| **middle** |  |  |  |  |
| HGS Tertiles (kg) |  |  |  |  |
| lower | -1.62 | -3.22~-0.02 | -2.00 | 0.047 |
| intermediate | 0.19 | -1.22~1.61 | 0.27 | 0.788 |
| upper | ref. |  |  |  |
| **high** |  |  |  |  |
| HGS Tertiles (kg) |  |  |  |  |
| lower | -0.96 | -2.39~0.47 | -1.32 | 0.186 |
| intermediate | -0.62 | -1.81~0.58 | -1.02 | 0.309 |
| upper | ref. |  |  |  |
| **Stratification by CLJ\*\*** |  |  |  |  |
| **low** |  |  |  |  |
| HGS Tertiles<sup>a</sup> (kg) |  |  |  |  |
| lower | -3.72 | -6.92~-0.53 | -2.32 | 0.023 |
| intermediate | -2.34 | -5.11~0.43 | -1.68 | 0.097 |
| upper | ref. |  |  |  |
| **middle** |  |  |  |  |
| HGS Tertiles (kg) |  |  |  |  |
| lower | -2.17 | -3.31~-1.04 | -3.76 | <0.001 |
| intermediate | -0.06 | -1.12~1.00 | -0.11 | 0.913 |
| upper | ref. |  |  |  |
| **high** |  |  |  |  |
| HGS Tertiles (kg) |  |  |  |  |
| lower | -1.30 | -2.78~0.19 | -1.72 | 0.086 |
| intermediate | -0.09 | -1.34~1.17 | -0.14 | 0.892 |
| upper | ref. |  |  |  |

**Abbreviations:** MMSE, Mini-mental state examination; *CI*, confidence interval; HGS, hand grip strength (kg); EDU, education; CLJ, cognitive level of the job; ref., reference.
\* Adjustment for age, sex, residence, marital status, smoking, alcohol consumption, physical exercise, BMI, total cholesterol, triglycerides, hypertension, diabetes, coronary heart disease, depressive symptoms, CLJ, and CLA.
\*\* Adjustment for age, sex, residence, marital status, smoking, alcohol consumption, physical exercise, BMI, total cholesterol, triglycerides, hypertension, diabetes, coronary heart disease, depressive symptoms, EDU, and CLA.

In addition, table 4 showed that the association between lower HGS and poor cognitive function was not statistically significant in elderly with high level of CLJ (*β*=*β* [95%CI]: −1.30 [−2.78~0.19], *P=*0.086), and less pronounced in individuals with middle level of CLJ (*β*=*β* [95%CI]: −2.17 [−3.31~−1.04], *P*<0.001) than those with low level of CLJ (*β*=*β* [95%CI]: −3.72 [−6.92~−0.53], *P*=0.023).

## 4. Discussion

Given the inconsistent results about the association between HGS and cognitive function in previous studies[3–5, 8–10], our study replicated and expanded previous findings with a strict control of confounding factors including sociodemographic factors, health behavioral factors, CV risk factors, and markers of cognitive reserve. In line with our expectation, the lower tertiles of HGS was significantly associated with lower cognitive function even after adjustment for confounding factors. Our foundings are consistent with prior studies [3–5,26,27], further corroborating the role of lower HGS in cognitive impairment in old age. HGS is considered as an effective and reliable measure of overall muscle strength [28], which may be an early marker of a generalized decrease in nervous system processing with age [3,29,30].

Consistent with previous studies, lower level of EDU and CLJ were related to decreased MMSE scores, suggesting high level of EDU and CLJ may attenuate cognitive decline in old adults. Although our study was cross sectional, it is in fact retrospective in nature as both EDU and CLJ happened in the early or middle-life stages. Furthermore, EDU and CLJ are two of important events in one’s life, information bias from self report may be minimal. Thus, a longitudinal relationship between EDU/CLJ and cognitive impairment is supported. Our results confirmed the conceptual view that various cognitive stimulations at the early and middle life stages may help to build cognitive reserve, thereby being related to better cognitive function in old age [14,16,19,20]. Participation in late-life leisure activity was also considered as a cognitive stimulation and may have beneficial effects on cognitive function [15,21,22]. Contrary to our expectation, no significant association was found between CLA and MMSE scores in our study. CLA has a broad spectrum (containing mental stimulation, social engagement, and physical activity, etc.) and its measurement has been hampered by lack of uniform definitions. In our study, CLA was defined as playing cards, scrabble, using computer, and reading, which may have a poor representation and discrimination. Further study is needed to assess the association between CLA and cognitive function.

Our main goal is to test whether cognitive reserve attenuate the relationship of lower HGS with cognitive function. We found that the correlation between lower tertiles of HGS and poor cognitive function was gradually attenuated with increase of level of EDU or CLJ and its statistical significance disappeared in individuals with high level of EDU or CLJ. To our knowledge, this is the first report on the role of cognitive reserve in the association of HGS with cognitive function. With the combination of other studies on the protective role of EDU or CLJ against cognitive impairment [14–17], our results supported the hypothesis that cognitive stimulation in the early or middle-life stages could help building up cognitive reserve, so as to buffer cognitive decline from risk factors, such as lower HGS.

Although our results confirm the relationship between HGS and cognitive function and the role of cognitive reserve, we are aware that the present study has some limitations. The first limitation may be represented by the present cross-sectional data, which does not take into account the causality question whether the decline in cognitive function is the consequence of reduction in hand grip strength or vice versa. Therefore, longitudinal studies are needed to further investigate the present observations before making suggestions. Another limitation was that although we were able to adjust for several key confounders, we cannot rule out the presence of unmeasured confounding. For example, we could not adjust for medication use factors, cardiac parameters, and arterial parameters parameters [31, 32].

## 5. Conclusions

In conclusion, we found that the lower HGS is associated with poor cognitive function in older age even after adjustment potential confounding factors and this relation was gradually attenuated with increase of level of EDU or CLJ. Our findings highlight the importance of maintaining HGS in delaying the onset of cognitive impairment. More importantly, cognitively stimulating activities in the early or middle-life stages should be enhanced to increase cognitive reserve and attenuate cognitive impairment from risk factors such as decrease of HGS. Future cohort or experimental studies are needed to clarify the causal relationships and to evaluate the effect of HGS maintaining and cognitive stimulations on delaying cognitive decline.

## Acknowledgements

The authors are grateful to all those who contributed to this study, including the research group of *Elderly Health and Modified Factors*, the Lu’an Center for Disease Control and Prevention, Fuyang Center for Disease Control and Prevention, Chengbei Township Health Center, Beishi community health service center, and jingjiu community health service center.

## Author Contributions

Conceptualization: Rong Wei, Kai-Yong Liu, Lin-Sheng Yang.

Data curation: Rong Wei, Lin-Sheng Yang.

Formal analysis: Rong Wei, Kai-Yong Liu.

Funding acquisition: Kai-Yong Liu, Fang-biao Tao, Lin-Sheng Yang.

Investigation: Rong Wei, Kai-yong Liu, Fang-biao Tao, Pei-ru Xu, Bei-jing Cheng,

Liang Sun, Qu-nan Wang, Qiang-wei Feng, Xiu-de Li, Lin-sheng Yang.

Methodology: Rong Wei, Kai-Yong Liu, Fang-biao Tao, Lin-Sheng Yang.

Project administration: Rong Wei, Fang-biao Tao, Lin-Sheng Yang.

Resources: Kai-Yong Liu, Fang-biao Tao, Lin-Sheng Yang.

Supervision: Rong Wei, Fang-biao Tao, Lin-Sheng Yang.

Validation: Rong Wei, Kai-yong Liu, Fang-biao Tao, Pei-ru Xu, Bei-jing Cheng,

Liang Sun, Qu-nan Wang, Qiang-wei Feng, Xiu-de Li, Lin-sheng Yang.

Visualization: Rong Wei.

Writing – original draft: Rong Wei, Kai-Yong Liu.

Writing – review & editing: Rong Wei, Kai-yong Liu, Fang-biao Tao, Pei-ru Xu, Bei-jing Cheng, Liang Sun, Qu-nan Wang, Qiang-wei Feng, Xiu-de Li, Lin-sheng Yang.

